# Frequency-Specific Resting-State MEG Network Characteristics of Tinnitus Patients Revealed by Graph Learning

**DOI:** 10.1101/2025.03.10.642147

**Authors:** Payam S. Shabestari, Harry H. Behjat, Dimitri Van De Ville, Christopher R. Cederroth, Niklas K. Edvall, Adrian Naas, Tobias Kleinjung, Patrick Neff

**Author notes:** Corresponding author *Email address:* (Payam S. Shabestari).

## Abstract

Tinnitus, the perception of sound without an external source, affects a significant portion of the population, yet its impact on brain communication diagram known as the functional connectome, remains limited. Traditional functional connectivity (FC) methods, such as Pearson correlation, phase lag index and coherence rely on pairwise comparisons and are therefore limited in providing a holistic encoding of FC. Here, we employ an alternative approach to estimate the entire connectivity structure by analyzing all time-courses simultaneously. This approach is robust even for short-duration recordings, facilitating faster functional connectome identification and real-time applications. Using resting-state MEG recordings from controls and individuals with tinnitus, we demonstrated that the learned connectomes outperform correlation-based connectomes in fingerprinting, that is, identifying an individual from test-retest acquisitions. Group-level analysis revealed distinct altered FC in tinnitus across multiple frequency bands, affecting the default mode, auditory, visual, and salience networks, suggesting a reorganization of these large-scale networks beyond auditory areas. Our study reveals that tinnitus presents highly individualized and heterogeneous whole-brain connectome profiles, highlighting the need to focus on individual variability rather than group-level differences to gain a more nuanced understanding of tinnitus. Personalized FC could enable patient-specific tinnitus models, optimizing treatment strategies for individualized care.

## 1. Introduction

Chronic subjective tinnitus is characterized by the ongoing conscious perception of sound without external acoustic sources [1], which can develop into a more complex condition known as “tinnitus disorder” associated with high levels of distress [2]. Affecting approximately 14.4% of the global population, tinnitus is common and often accompanied by severe burdens [3], including comorbidities such as depression and anxiety. Tinnitus is typically considered to result from noise trauma and/or hearing loss, leading to a lack of peripheral auditory input and subsequent maladaptive changes in the auditory pathway and central nervous system, which are believed to cause the perception of phantom sounds [4]. These pathological changes are reflected in distinctive brain activity patterns, with increased gamma and delta, and decreased alpha band activity in the auditory cortical regions, as observed by EEG and *MEG studies [5, 6]. Additionally, tinnitus-related changes have been observed in both global and modality-specific functional networks, which typically show increased connectivity within and between the auditory network, the default mode network, attention networks, and the visual network [7, 8, 9]*.

Complex brain functions and neuropathologies such as tinnitus are supported by anatomically and functionally interconnected networks of brain regions [10, 11, 12, 13]. Functional connectivity (FC) is the most common technique to identify these networks through fMRI [14] or M/EEG recordings [15]. In this framework, brain regions are depicted as vertices, while statistical metrics of dependency like Pearson correlation, Phase lag index [16], or methods relying on time-resolved evaluations of cross and power spectral densities such as Coherence between regional time-courses, serve as the weights of edges. However, despite its widespread use, FC has two major short-comings that undermine the reliability and interpretability of the networks it recovers. First, FC assumes that brain regions are functionally connected if their activities are temporally correlated. However, this correlation can occur even if the regions do not actually exchange functional information. For example, misleading correlations can arise from noise that is temporally synchronized due to physiological factors such as respiration, heart rate, and movement [17]. Furthermore, the magnitude of spurious correlations is influenced by the signal-to-noise ratio (SNR) of the brain areas involved. Regions with low SNR are particularly susceptible to these erroneous correlations caused by temporally correlated noise. Secondly, functional connectomes inherently offer only a pairwise view of brain dynamics, simplifying the brain into dyads. This simplicity, restricts the exploration of unique features within human brain networks.

Finn et al. [18] demonstrated that certain FC edge weights are idiosyncratic—both distinctive to each individual and consistent across repeated fMRI sessions. Crucially, the idiosyncratic edges that aid in identifying individuals were also found to be the most predictive of behavioral traits [18]. These discoveries have inspired numerous studies emphasizing idiosyncratic rather than generalizable FC patterns, leading to the emerging field of brain fingerprinting. Recently, few studies have started to explore connectome fingerprinting across different functional neuroimaging modalities, such as MEG [19], expanding the applicability of this approach beyond fMRI.

Using principles from the recently emerged field of graph signal processing [20, 21], and motivated by its promising applications to neuroimaging data [22, 23, 24, 25], in this work we used an alternative approach to define subject-specific, sparse functional networks [26]. The underlying assumption for this method is that brain activity should be manifested as a smooth signal residing on an underlying manifold, which can be represented in discrete form as a graph/network [26, 27]. Intuitively, a signal is considered smooth on a graph if neighboring brain regions—those connected by edges with strong weights—exhibit similar activity levels. This reflects the idea that functionally or anatomically connected areas in the brain are likely to show correlated activity patterns. Smoothness is typically quantified by measuring how much the activity values differ between connected regions; smaller differences across strongly connected nodes indicate a higher degree of smoothness. Mathematically, this concept is formalized using the graph Laplacian, which allows us to compute an overall smoothness score that captures how well the observed brain activity conforms to the structure of the network. Prior work on EEG [28, 29] and fMRI [30, 31] data has shown the applicability of this approach in revealing individualized brain networks with application to motor imagery decoding and subject identification. This approach offers two key advantages: i) it mitigates misleading correlations caused by temporally synchronized noise, as graph learning treats each time point as an independent observation and ii) by leveraging all time courses simultaneously, it addresses limitations of dyadic networks inherent in pairwise correlation methods, providing a more robust and holistic understanding of functional brain connectivity.

We further investigated several aspects of brain connectivity in the context of tinnitus: i) The robustness of the generated graphs when constructed using only a small fraction of the available time samples, demonstrating the method’s efficiency in data-limited scenarios. ii) A statistical group-level comparison between healthy controls and tinnitus individuals, which highlighted significant differences in connections between brain regions associated with auditory processing and memory. iii) The examination of within-session brain connectivity fingerprints, showcasing the ability to identify an individual’s functional connectivity (FC) profile within a population, emphasizing the consistency of brain connectivity patterns within individuals. iv) The analysis of the distribution of connections with the highest fingerprint, which allowed for subject identification. This revealed significant involvement of brain regions within key functional networks in tinnitus, namely medial posterior cingulate and ventro-medial prefrontal cortex complexes. The aim of this study was to investigate whether whole-brain functional connectivity patterns are reliably stable across both healthy individuals and tinnitus patients. We aimed to assess whether functional connectomes could be used to identify individual patients, and to examine potential alterations in the spatial organization of brain networks in tinnitus patients, suggesting unique changes in their functional connectivity patterns.

## 2. Materials and methods

### Study population

The research received ethical approval from the Regional Ethics Committee in Stockholm, *Regionala* etikprövningsnämnden (Dnr:2019-05226). All participants provided written informed consent after being fully briefed on the study’s purpose, scope, and potential risks. The study strictly adhered to the ethical principles of the Declaration of Helsinki. Control individuals consisted of 26 individuals with normal hearing that were recruited via the online platform (accindi.se). Participants with chronic and constant tinnitus (tinnitus always present in silence for > 1 year) were recruited via the accindi platform (n = 8), the Swedish Tinnitus Outreach Project (STOP; n = 4) and the Karolinska Hospital (n = 8). Five participants were excluded due to technical difficulties (MR), claustrophobia and/or low quality signal resulting in a final group size of n = 23 for control group and n = 18 for tinnitus group. Exclusion criteria included pregnancy, sound sensitivity (hyperacusis), psychiatric or neurological conditions, drug use, and non-removable metal implants. Control participants self-reported their sex, resulting in a balanced distribution of males and females, and one identifying as other. Their mean age was 28.4 years (SD ± 5.8), with 88% being right-handed. Tinnitus participants comprised 7 females and 11 males. Their mean age was 36.8 years (SD ± 7.7), with 94.4% being right-handed. we included all available participants and therefore did not enforce strict demographic matching. However, to assess potential confounding effects, we conducted a post-hoc demographic comparison using nearest neighbor matching (matchit R function, MatchIt package). The distribution of sex was relatively balanced and did not change substantially before and after matching. Although a significant difference in age remained between the tinnitus and control groups (ranging from approximately 28 to 36 years), we note that this range falls within a period of stable brain maturation, where major developmental changes are not typically expected. Therefore, we believe it is unlikely that this moderate age difference substantially influenced our connectivity findings (see Table 1).

**Table 1:**
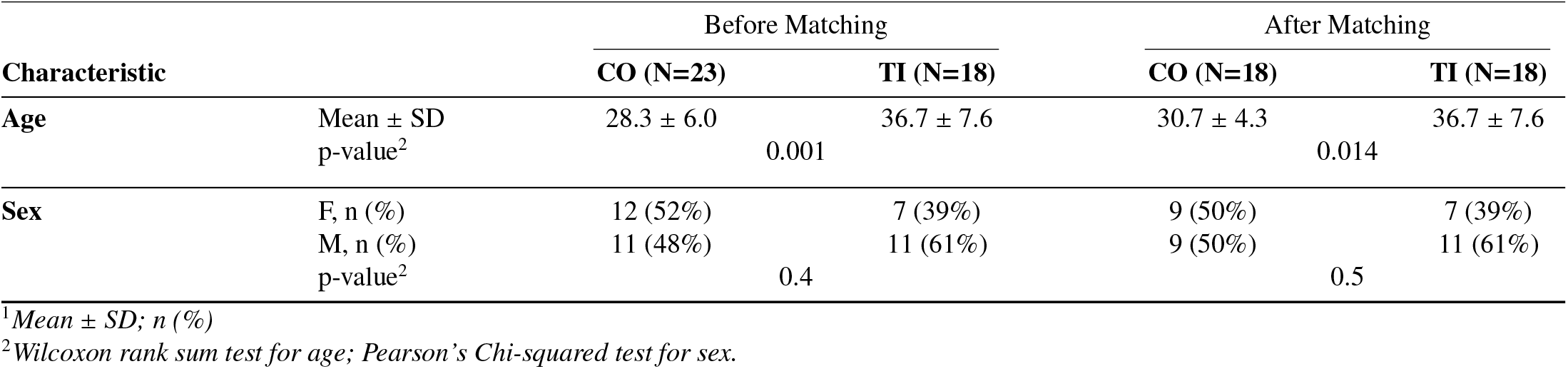
Statistical comparison of the groups (controls vs tinnitus) before and after matching.

Participants attended the national magnetoencephalography facility (NatMEG) for a single session, where they received detailed procedural information, confirmed the absence or presence of tinnitus or hearing issues, and completed an online survey assessing stress, anxiety, depression, and hyperacusis before undergoing auditory assessments. Resting-state MEG recordings were conducted, followed by a structural MRI scan for each participant using a 3 Tesla GE MR750 Discovery scanner at the MR Center, Karolinska Institutet.

### MEG and MRI data acquisition

The MEG data consisted of recordings from 18 individuals with tinnitus and 23 controls acquired using a 306-channel Elekta Neuromag184 TRIUX system. Standard MEG preparation procedures were implemented, which included the placement of position indicators (cHPI coils) to monitor movement, 3D registration of the scalp using Polhemus for alignment with structural MRI, and the attachment of electrodes to mitigate muscle artifacts from heart and eye movements. Participants were instructed to sit in a relaxed manner and watch a nature film devoid of sound displayed on a screen in front of them. The silent nature movie was projected onto a screen measuring 72 × 44 cm using an FL35 LED DLP projector positioned outside the magnetically shielded room. The silent movie was used as a neurophysiological “reset” to help participants reach a stable, neutral mental state before the resting-state MEG recording. This minimized variability from prior cognitive or emotional states and reduced anticipatory arousal. It also promoted consistency across participants by establishing a comparable baseline. Importantly, the silent format avoided auditory or language-related stimulation that could affect resting-state activity. The MEG session concluded with a 5-minute resting state recording, during which participants were instructed to maintain a relaxed posture and continue watching the movie as they had done previously during the measurement.

The MRI data consisted of 3D T1-weighted magnetization-prepared rapid gradient-echo (MPRAGE) sequence structural images. These images were acquired using a GE Discovery 3.0T MR scanner, with a voxel size of 1x1x1 mm, a field of view of 256 mm, repetition time (RT) of 2300 ms, and echo time (ET) of 2.98 ms.

### MEG and MRI data processing pipeline

The MEG data, sampled at 5000 Hz, underwent several preprocessing steps. Signal-Space Projection (SSP) vectors [32] were derived from empty-room recordings conducted before and after the experiment to filter out environmental noise from sources external to the subject and the MEG system. Continuous head position indicator (cHPI) coil signals were used to estimate and track head movements for compensation during recording (see Supplementary Information for a comparison of head movement between the two groups). Automatic identification and correction of noisy and flat MEG channels, along with crosstalk compensation, were performed using spatiotemporal Signal-Space Separation (tSSS) in 10-second intervals [33, 34]. After downsampling to 250 Hz and applying a low-pass filter to prevent aliasing, bandpass filtering between 0.1 and 80 Hz was applied. Independent Component Analysis (ICA) was employed to decompose the data for artifact correction [35]. The minimum number of components capturing at least 95% of data variance was determined. Specifically, for ECG artifacts, a bandpass filter was applied, and relevant components were removed using the Cross-Trial Phase Statistics method [36]. Muscle artifacts were identified and eliminated using the methodology outlined in [37], which involves identifying components with a positive slope in the power spectrum between 7–75 Hz, a peripheral or non-central topographic focus, and low spatial smoothness (i.e., a single, sharply localized activation rather than widespread neural patterns). The signals were bandpass filtered into standard frequency bands: Delta (0.5–4 Hz), Theta (4–8 Hz), Alpha (8–13 Hz), Beta (15–30 Hz), and Gamma (30–80 Hz). This filtering was used to estimate brain activity within each frequency range for further analysis.

We conducted anatomical cortical surface reconstructions of the MRIs using `FreeSurfer` software version 7 [38]. This processing involved cortical surface reconstruction and the generation of source space models. Subsequently, we created Boundary Element Model (BEM) surfaces, encompassing the inner skull, outer skull, and outer skin (scalp), employing the watershed algorithm [34, 39]. A surface-based source space was established, representing bilateral hemispheres, and recursively divided with octahedron spacing. For each subject, a BEM model and its corresponding solution were constructed using the linear collocation method [34]. To ensure accurate alignment, co-registration between the MRI and the head model was performed, initially using three fiducial points provided in the MRI. This alignment was refined through 40 iterations of the Iterative Closest Point (ICP) algorithm [40], with outlier points exceeding a distance of 5 cm excluded and the fitting process reiterated. For each subject, the forward solution was computed utilizing the source model, BEM model, and co-registration data. We computed the noise covariance of the recordings based on empty room recordings. Estimation of noise covariance was carried out using both empirical [34] and shrunk [41] methods, with the best estimator chosen based on log-likelihood and cross-validation with unseen data [42]. Applying the linear minimum-norm inverse method (dSPM) with the noise covariance and forward solution enabled determination of the inverse solution [34], yielding source time courses for each vertex in the source space. Subsequently, a single time course was generated for each brain label by averaging source time courses within vertices located in that specific brain label, irrespective of their orientation. Brain labels and cortical parcellation were derived from the Desikan-Killiany Atlas [43], comprising 68 bi-hemispheric parcels. All MEG processing analysis was performed using `MNE` software (Version 1.6.1).

### Construction of functional networks

Let 𝒢 = (𝒱, ℰ, **A**) denote a weighted, undirected graph, where 𝒱 = {1, 2, …, *N*} represents the graph’s finite set of *N* vertices (nodes), ℰ denotes the graph’s edge set, i.e., pairs (*i, j*) where *i, j* ∈ *ε*, and **A** denotes the graphs’ adjacency matrix, which is symmetric with elements *A*_*i*, *j*_ = *A*_*j,i*_ representing the weight of (*i, j*). The weights in the adjacency matrix indicate the strength of the connection, or similarity between two corresponding vertices, therefore, *A*_*i*, *j*_ = 0 if there is no connection/similarity between vertices *i* and *j*. In this work we consider graphs with no self-loops, i.e., *A*_*i,i*_ = 0. The graph’s combinatorial Laplacian matrix is defined as:

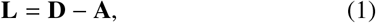

where **D** is the diagonal matrix of vertex degrees with its elements given as

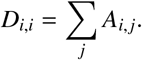

Let **f** ∈ ℝ^*N*^ denote a graph signal, that is, a real signal defined on the vertices of 𝒢whose *n*-th component (**f**[*n*]) represents the signal value at the *n*-th vertex of 𝒢. The total variation (TV) of a graph signal **f** on graph 𝒢 can be quantified using **L** as [44]:

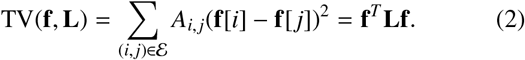

Larger values of TV(**f, L**) indicate greater changes of **f** on 𝒢, i.e., higher spatial variability, and thus, lower spatial smoothness. Using this notion of smoothness, a sparse graph structure can be inferred from a given set of observations [27]. Let

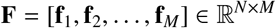

denote a matrix that stores a set of *M* measurements on a domain with *N* elements, and let **Z** denote an *N*×*N* matrix with entries that represent the Euclidean distance (or any other distance metric) between pairs of rows in **F**, i.e.,

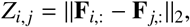

where **F**_*k*,:_ denotes the *k*-th row of **F**, that is measurements from the *k*-th element. A graph structure can be inferred from **F** via the optimization [26]:

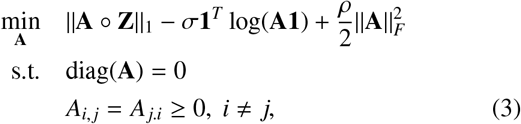

where ∘ and∥·∥_*F*_ denote the Hadamard product and the Frobenius norm, respectively. σ and ρ are regularization parameters; intuitively, smaller values of ρ yield sparser graphs by penalizing edges between vertices with larger *Z*_*i*, *j*_ [26]. The first term in (3) enforces smoothness by invoking that [26]:

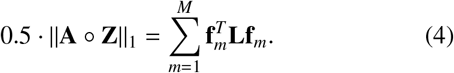

Smooth graph signals reside on strongly-connected vertices, thus, the vertices are expected to have smaller distances. The second term enforces degrees to be positive and improves overall connectivity.

To evaluate the performance of the constructed FC graphs, we used the Phase Lag Index (PLI) method. PLI is a reliable measure of phase synchronization that remains unaffected by common sources, such as volume conduction or active reference electrodes. This method is widely utilized to create brain FC networks and it helps in determining the phase relationship between two signals, indicating the extent to which signal leads or lags behind another [16]. Each element of the PLI matrix (*N* × *N*) can be quantified as:

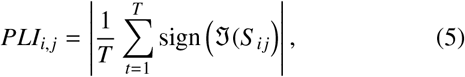

where *T* denotes the total number of time-points and ℑ (*S* _*i j*_) denotes the imaginary component of the averaged cross-spectral density (CSD) across frequency bins. The CSD is computed between time-courses of two cortical regions *i* and *j* (i.e. **F**_*i*,:_ and **F** _*j*,:_) using Morlet wavelets, and it has the dimensions of the number of time points by the number of frequency bins. *PLI*_*i*, *j*_ = 0, means that signal **F**_*i*,:_ leads and lags signal **F** _*j*,:_ equally often, while a value greater than 0 means that there is an imbalance in the likelihood for signal **F**_*i*,:_ to be leading or lagging. A value of 1 means that signal **F**_*i*,:_ only leads or only lags signal **F** _*j*,:_.

### Network robustness

To assess the stability of the learned graphs across varying numbers of samples and learning parameters, we followed these steps: We constructed graphs using four different σ and ρ regularization parameters chosen from 0.25, 0.5, 1, and 1.5. These graphs were created for different numbers of samples ranging from 2 to 300 seconds. For each selected number of samples, we randomly sampled data points from the entire recording, repeating this process five times per number of samples. For each parameter configuration, we designated the graph constructed using all samples (300 seconds) as the ideal graph (*G*_*ideal*_). We then compared each graph to the ideal graph by computing the Portrait divergence (PDiv) and Euclidean distances between them. The Portrait divergence compares the “graph portraits” which are distributions of shortest path lengths between nodes and was calculated using the method described in [45] and the Euclidean distance was computed by subtracting the upper triangles of the adjacency matrices and taking the 2-norm of the resulting differences. Figure 1.a presents an overview of the processing steps, where graph structures are inferred from brain parcel signals across various time windows and compared to the optimal graph, which is constructed using the entire recording.

**Figure 1.**
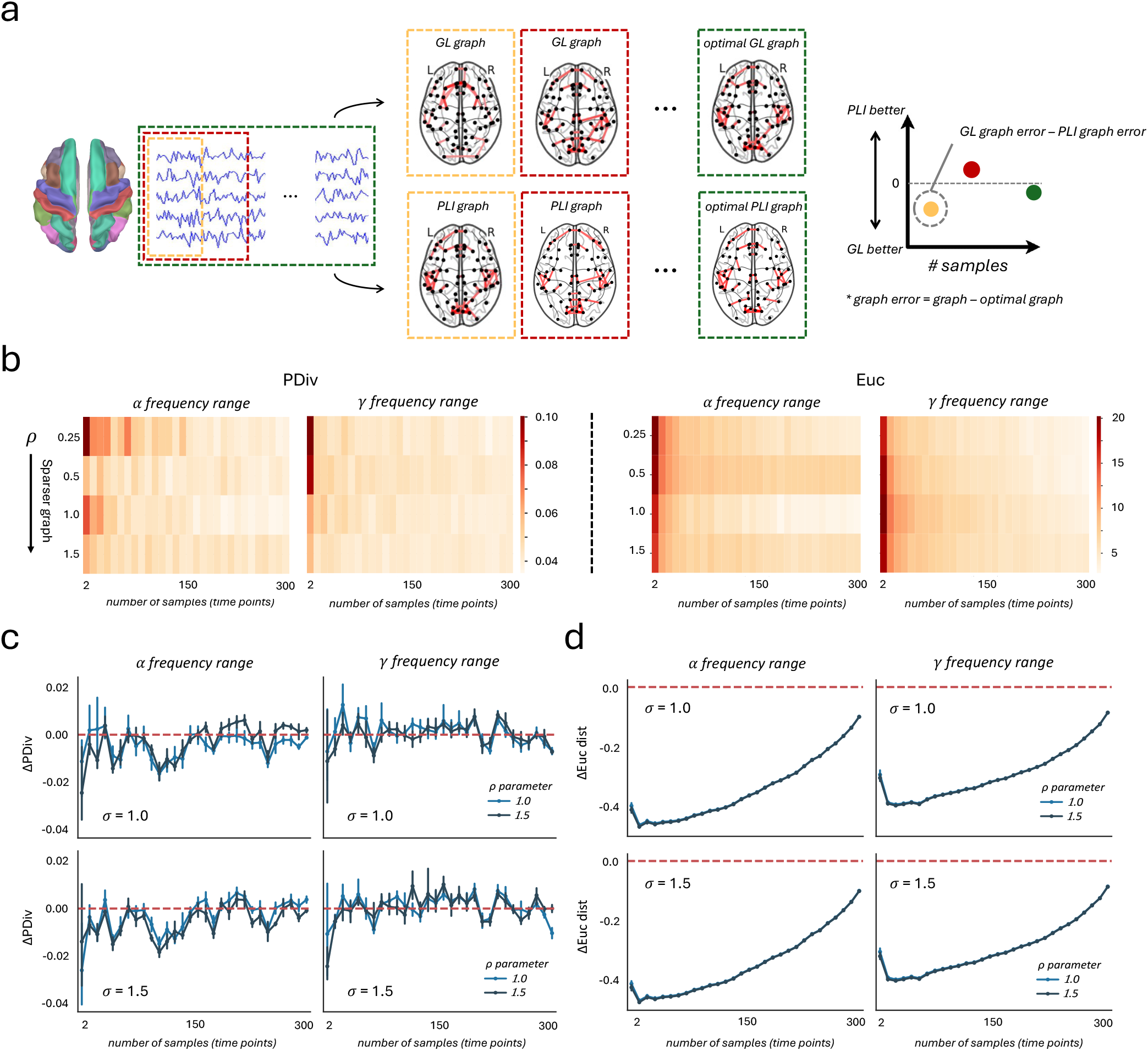
(**a**) Overview of processing steps: graph structures inferred from brain parcel signals across various time windows, and compared to the optimal graph learned from the entire recording. The same analysis was conducted using the PLI method, and the results from both approaches were then compared. (**b**) PDiv and Euclidean distances were calculated between each constructed graph and the optimal graph at alpha and gamma frequency ranges. Differences in PDivs (**c**) and Euclidean distances (**d**) between PLI and GL methods; negative values indicate that the GL method results in less divergence (greater similarity) to the optimal graph compared to the PLI method.

### Statistical group comparison

To compare group-level global cortical connectivity between controls and tinnitus patients, we first normalized each FC using its Frobenius norm. Edges with weights below 1% of the maximum edge weight were excluded from the analysis. Statistical tests were conducted as two-sided, with normality evaluated via the Shapiro-Wilk test (stats.shapiro Python function, scipy package). For data not meeting the normality assumption, the non-parametric Mann-Whitney U test (mannwhitneyu Python function, scipy package) was employed. Bonferroni correction was applied to control for multiple comparisons across all 2,278 edges between the 68 brain regions, and edges with p-values below 0.05 were considered statistically significant.

### Subject identifiability of functional networks

The fundamental concept of subject identifiability revolves around assessing whether the connectivity profiles, or graph structures, corresponding to the same individual demonstrate more similarity compared to those across different individuals. We employed identifiability measures from [46] to construct an *S*× *S* identifiability matrix **M** for the given FC matrices. In this context, *S* represents the number of subjects, and each element *M*_*k,l*_ is the Pearson’s correlation between 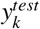 and 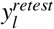, where 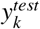 and 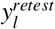 denote the test (first half) and retest (second half) feature vectors of subjects *k* and *l*, respectively. First, for each subject—within any of the groups—we generated two graphs: i.e. test and retest (each 2.5 minutes). In these graphs, vertices denote the center of mass for macro-scale cortical regions, as defined by the Desikan-Killiany atlas [43], encompassing a total of 68 regions. The graph structure associated to each test or retest was inferred via an optimization outlined in Eq. 3. Given that the learned graphs are symmetric, we only used the upper triangle part of their adjacency matrices to compute the identifiability matrix. Finally, we sorted the correlation values for each subject, determining the rank of correlation of the same subject between the test and retest FCs. Identification ranks offer a reliable group-level estimate of identifiability at the individual connectome level, with lower values indicating better individual identifiability. To further investigate the observed heterogeneity, we applied spectral clustering to the symmetric similarity matrix derived from the identifiability matrix. This similarity matrix was computed by calculating the cosine similarity between all pairs of rows in the identifiability matrix. Spectral clustering involves computing a low-dimensional embedding of this affinity matrix using the eigenvectors of its Laplacian, followed by clustering (cluster.SpectralClustering Python function, sklearn package) in the embedded space. We set the number of clusters to three, based on visual inspection and prior expectations about subgroup structure.

### Spatial specificity of functional network fingerprints

The spatial specificity of functional connectivity (FC) fingerprints was assessed using edgewise intra-class correlation (ICC) based on a one-way random effects model [47]:

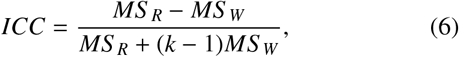

where, *MS* _*R*_ denotes the variance between subjects (mean square for rows), *MS* _*W*_ represents the residual variance, and *k* is the number of sessions. ICC quantifies the consistency between measurements and is widely used in reliability studies. Higher ICC values indicate greater consistency, reflecting that most of the variance originates from differences between individuals rather than measurement errors or test-retest variability. Conversely, lower ICC values suggest that random error or inconsistencies between test and retest contribute more significantly to the variance. In this work, ICC was applied, following previous studies, to measure FC stability between pairs of brain regions (edges) across test-retest sessions. A high ICC for an edge signifies that its connectivity remains highly consistent for a given individual across sessions and varies systematically across subjects, emphasizing the edge’s distinctiveness or “fingerprint”.

## 3. Results

### Network robustness

Figure 1.b illustrate the portrait divergence (PDiv) and the Euclidean distance, respectively, between each learned graph and the optimal graph across different configurations of the learning parameter ρ (σ is set to 1). As the number of time samples used for training increases, both the PDiv and Euclidean distance decrease, reflecting a closer alignment with the ideal graph. To further benchmark the learning method against the PLI approach, we analyzed both PDiv and Euclidean distance within connectomes obtained using the PLI method and computed the differences by subtracting the corresponding values derived from the graph learning (GL) method. Figures 1.c and d illustrate these differences for the alpha and gamma frequency bands across various σ and ρ configurations. Negative values for ΔPDiv (Figure 1.c) and ΔEuc (Figure 1.d) indicate that the GL method has a lower divergence and distance from the ideal graph. This suggests that the GL method achieves a greater similarity to the ideal graph in terms of both local and global structures compared to the connectomes derived using the PLI method.

### Statistical group comparison of functional networks of tinnitus and healthy controls

Group-level statistical comparisons of the graphs learned from the control and tinnitus groups revealed significant differences in functional connectivity in specific frequency bands: theta (4–8 Hz), alpha (8–13 Hz), and gamma (30–80 Hz) (Figure 2). No significant differences were observed in the delta (0.5–4 Hz) or beta (13–30 Hz) bands. In the theta band, significant connections were identified between the fusiform gyrus and the banks of the superior temporal sulcus in the right hemisphere (*t*(39) = 4.65, *p* = 0.042), the cuneus and the lateral occipital cortex in the left hemisphere (*t*(39) = 4.54, *p* = 0.044), and the paracentral-rh lobule and the precentral-lh gyrus (*t*(39) = 4.27, *p* = 0.046). These connections were notably weaker in individuals with tinnitus, suggesting reduced theta band connectivity in these regions. In the alpha band, significantly different connections were observed between the parahippocampus and the insula-lh (*t*(39) = 4.74, *p* = 0.039) and the superior frontal gyrus in both hemispheres (PHC-lh vs SFL-lh: *t*(39) = −5.42, *p* = 0.031, PHC-lh vs SFL-lh: *t*(39) =−5.39, *p* = 0.031). Additionally, the lateral orbitofrontal cortex showed stronger connectivity with the transverse temporal gyrus in individuals with tinnitus (*t*(39) = −3.68, *p* = 0.046). These findings suggest that tinnitus is associated with increased alpha band connectivity in regions related to memory and frontal processing. In the gamma band, stronger connections were observed in the control group between the superior frontal gyrus and the frontal pole (SFL-rh vs FP-rh: *t*(39) = 5.51, *p* = 0.035, SFL-lh vs FP-rh: *t*(39) = 4.88, *p* = 0.040, SFL-lh vs FP-lh: *t*(39) = 4.75, *p* = 0.041). In contrast, individuals with tinnitus exhibited stronger connectivity between the inferior parietal cortex and the precuneus (*t*(39) = 4.0, *p* = 0.045), the inferior parietal cortex and pars orbitalis (*t*(39) =−4.71, *p* = 0.041), the superior parietal cortex and the paracentral gyrus (*t*(39) = 3.47, *p* = 0.046), and the pars orbitalis and the banks of the superior temporal sulcus (*t*(39) = −6.03, *p* = 0.029). These patterns suggest a higher activity of the gamma band in patients with tinnitus in networks related to attention and sensory processing. The correlation between the functional connections that showed significant group-level differences and the pure tone audiometry (PTA) scores across participants was assessed and presented in supplementary Figure S5.

**Figure 2.**
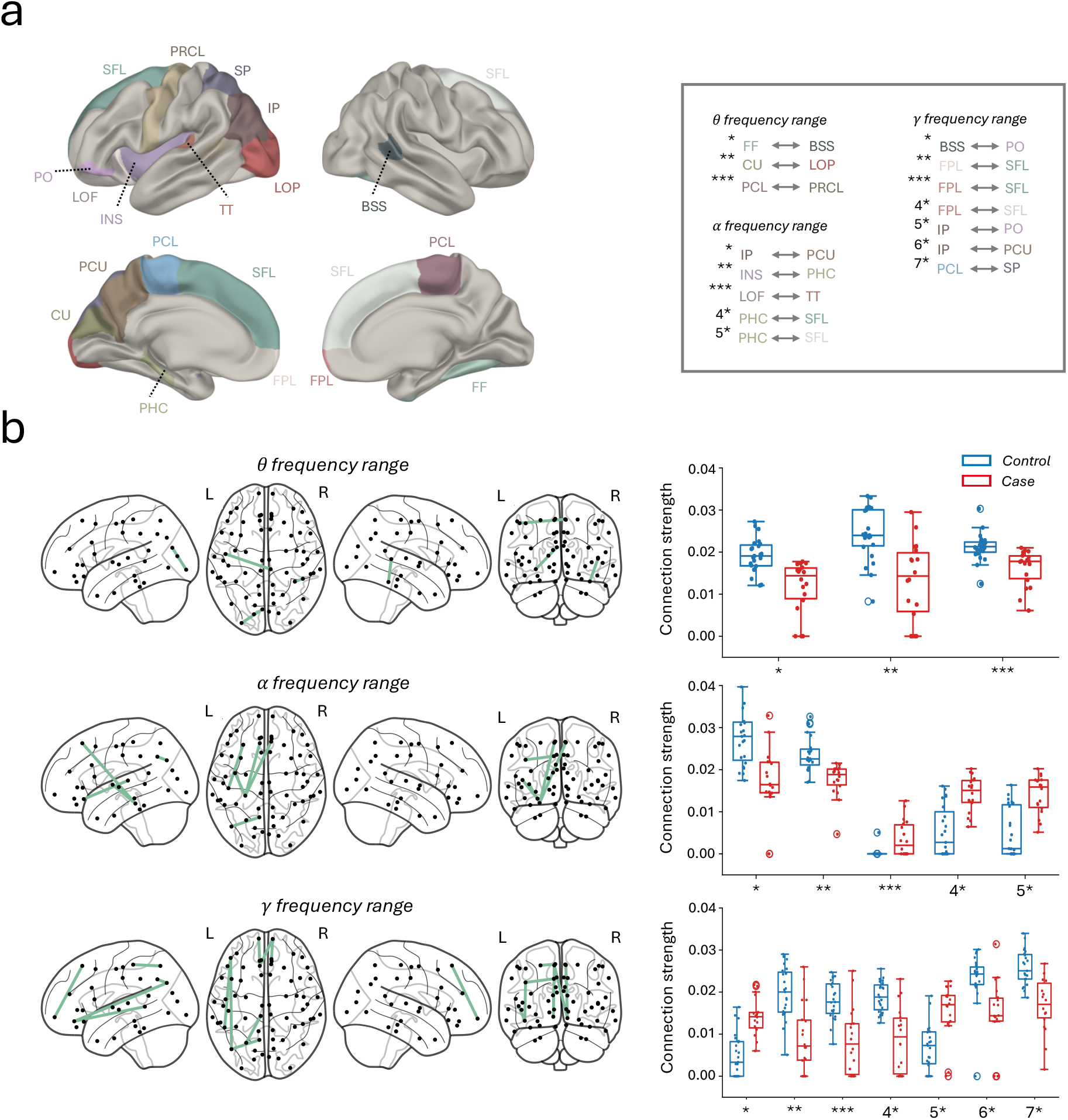
(**a**) The significantly different edges over ROIs (i.e., between centroids of the ROIs) are shown on a semi-inflated standard brain surface (left/right hemispheres, lateral and medial views). Abbreviations not specified above: FF fusiform gyrus, BSS banks of the superior temporal sulcus, CU cuneus, LOP lateral occipital cortex, PCL paracentral lobule, PRCL precentral gyrus, IP inferior parietalclobule, PCU precuneus, INS insula, PHC parahippocampal gyrus, LOF lateral orbitofrontal cortex, TT transverse temporal gyrus, SFL superior frontal gyrus, PO pars orbitalis and FPL frontal pole. (**b**) Significantly different graph edges (*p* < 0.05) between the two groups are illustrated for the canonical frequency bands (i.e theta, alpha and gamma). On the right, boxplots display the distribution of these connection strengths across subjects for each band in both control (blue) and tinnitus patient (red) groups. Connection IDs (x-axis labels) in the boxplots correspond to those shown in panel (a).

### Within-session functional network fingerprints

The identification rank of each individual using both the PLI and GL methods is presented in Figure 3.a for both groups in two frequency ranges: alpha and gamma (see Supplementary Figure S6 for comparisons with other common methods, including PLV and Coherence). Graph learning outperforms PLI, achieving first rank for the majority of subjects, suggesting that the learning approach better captures subject-specific idiosyncrasies in function. In this analysis, both the learning parameters σ and ρ were set to 1. We investigated within-session brain fingerprints in two independent groups for a total of N = 41 subjects (control group: N = 23 and tinnitus group: N = 18). Our approach can be summarized in three steps: (i) we first estimated the FCs of each subject during the first (test) and second halves (retest) of MEG acquisitions separately (see Section 2 for details). (ii) We then estimated the degree of within-session brain identification or “brain fingerprint” at the wholebrain level for each group separately, through a mathematical object called an identifiability matrix (Figure 3.a). The identifiability matrix provides two useful metrics for brain fingerprinting: the degree of similarity of each subject to himself (Iself, diagonal elements, Figure 3.b) and the degree of brain discriminability, conceptualized as the extent to which subjects were more similar to themselves than others (Idiff, section 2 for details) (density and boxplots in Figure 3.a). The fingerprinting results revealed significant idiosyncrasies in individual subject networks, particularly in tinnitus cases. (iii) we then applied spectral clustering with three clusters to the symmetric similarity matrix. For better visualization, the symmetric matrix was projected into a 2D space using Multidimensional Scaling (MDS) (manifold.MDS Python function, sklearn package). MDS models similarity or dissimilarity data as distances in a geometric space. Figure 4.a displays the MDS representation of individuals for the alpha and gamma frequency bands, with clustering results (control: blue, tinnitus: red). Marker sizes correspond to individuals’ *I*_*di f f*_ values, where larger marker sizes indicate greater brain discriminability from others. In contrast, Figure 4.b presents the MDS representation for both groups based on individual PLI networks. This analysis suggests that there is no discernible group clustering between the two groups.

**Figure 3.**
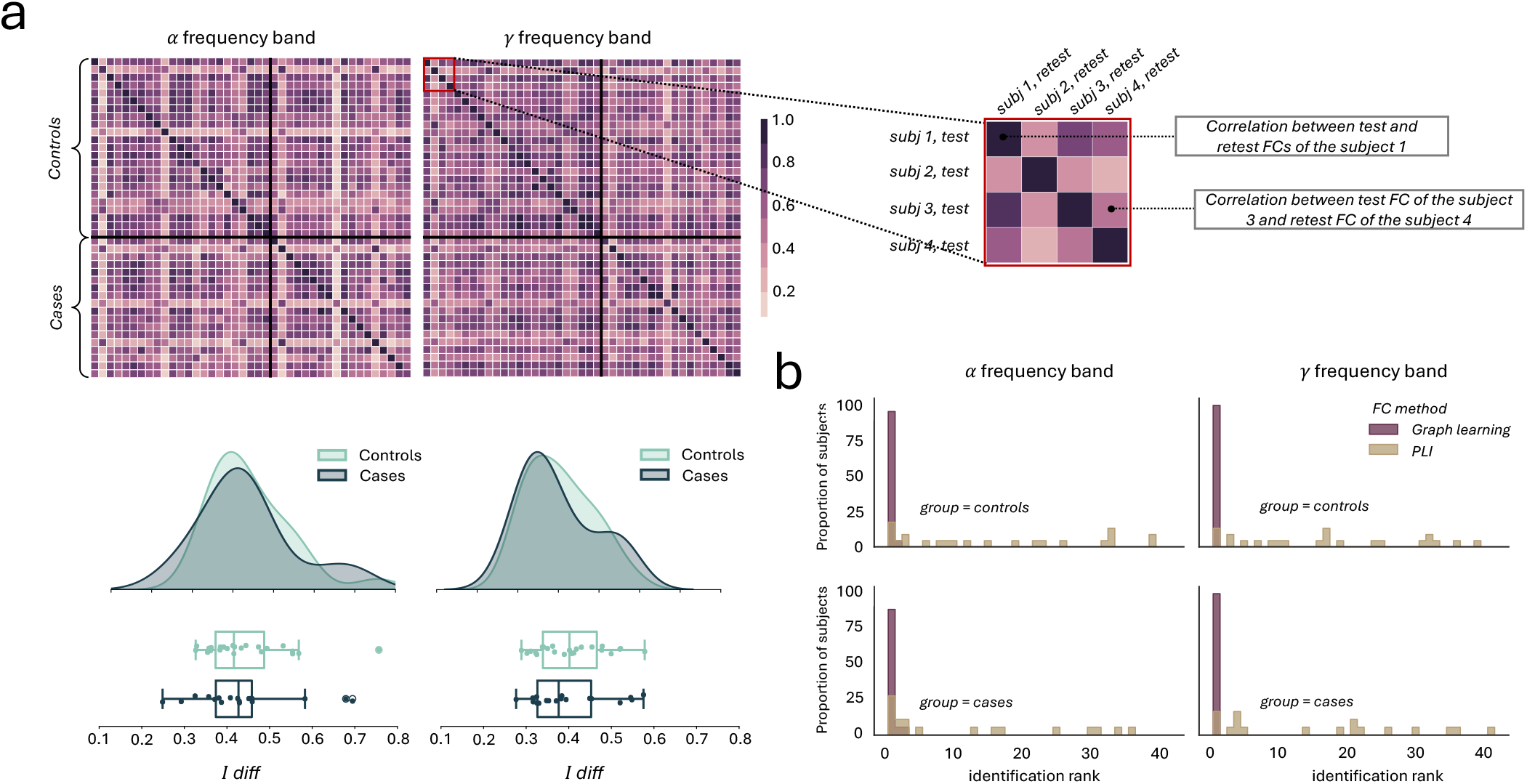
(**a**) The identifiability matrix shows within-subjects similarity (Iself, diagonal elements) and between-subjects similarity (off-diagonal elements) across all subjects (tinnitus + control) for frequency bands alpha and gamma. The density and box plots of Idiff values are shown below. The Idiff value for each subject is calculated as the difference between their Iself value and the average of their off-diagonal values, highlighting the subject’s distinctiveness in functional connectivity.(b) Bar plots showing the identification ranks in two groups (tinnitus and control) derived via the GL and PLI methods, comparing the test and retest FCs among subjects.

**Figure 4.**
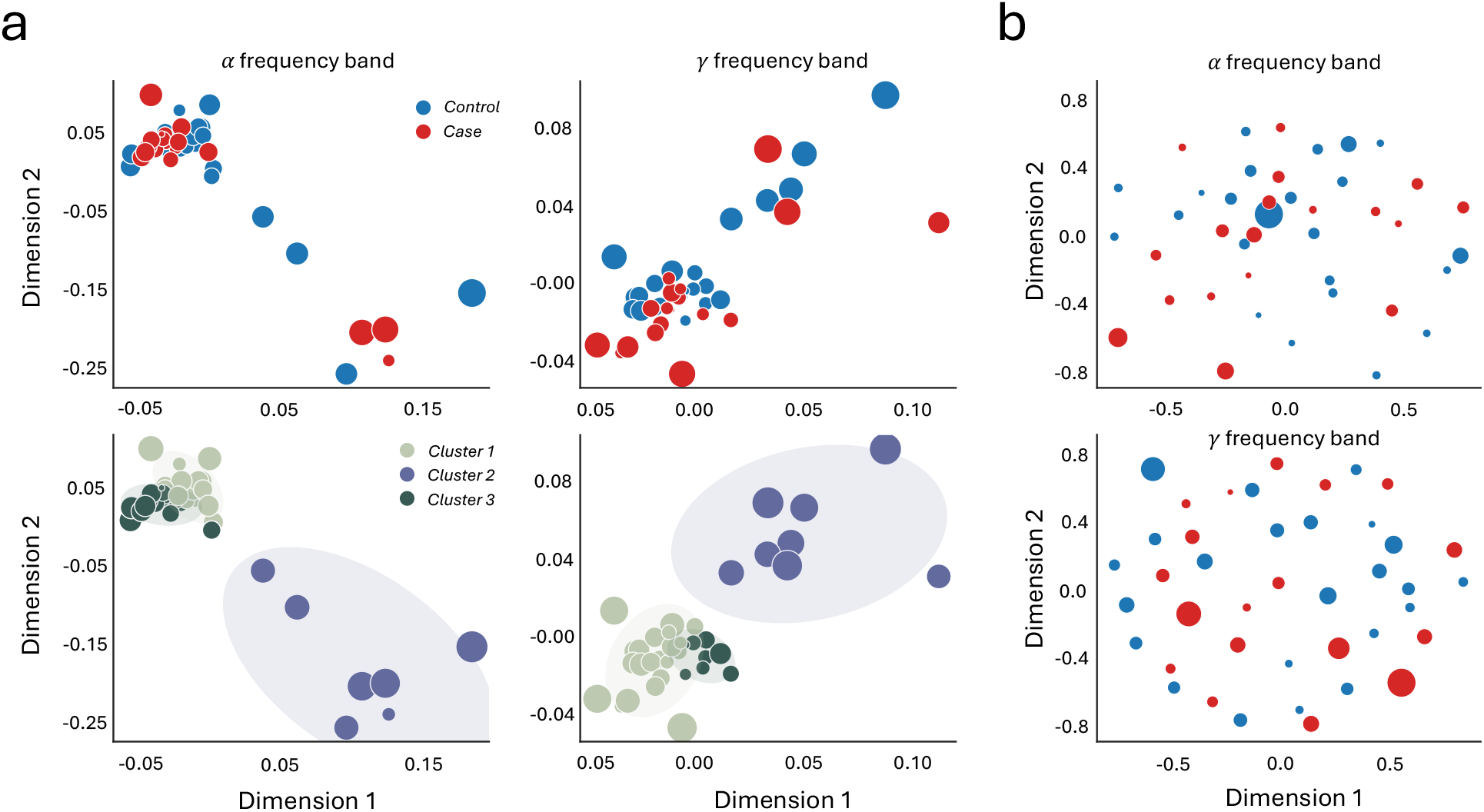
(**a**) The 2D MDS representation of the similarity matrix, derived from the identifiability matrix for the alpha and gamma frequency bands, shows a clear grouping of controls and tinnitus patients. Below this, clustering results into three distinct clusters further highlight a more robust group formation compared to the PLI-based fingerprinting as shown in (**b**). The silhouette scores for alpha and gamma band clustering are 0.210 and 0.202, respectively. In both panels, the marker sizes correspond to individuals Idiff values, where larger marker sizes indicate greater brain discriminability from others.

### Spatial specificity of functional network fingerprints in tinnitus

In these analyses, our goal was to identify brain regions (or nodes) whose functional connectivity with the rest of the brain contributed significantly to the observed variability within the tinnitus group. To achieve this, we evaluated the spatial specificity of brain fingerprints using edgewise intra-class correlation (ICC) (see Section 2 for details). This analysis revealed a spatial reconfiguration of the most identifiable edges in individuals with tinnitus, particularly in the alpha and gamma frequency ranges (Figure 5.a and b, left panels). To further investigate this spatial reconfiguration, we analyzed the pattern of ICC nodal strength for each brain region. The nodal strength was calculated as the sum of the ICC values for all edges connected to a given brain region. Regions with the highest nodal strength—specifically, those in the top 50th percentile—were visualized by rendering them onto the cortical surface (Figure 5.a and b, right panels). This approach highlights key regions contributing to altered functional connectivity in tinnitus. In the alpha band, spatially discriminative patterns were observed in the bilateral isthmus cingulate and the right precuneus—regions typically associated with the default mode network—as well as the left pericalcarine gyrus. In the gamma band, altered spatial configuration involved the left frontal pole, right pericalcarine gyrus, and left medial orbitofrontal cortex. (see supplementary for spatial configuration of brain fingerprints in control group.)

**Figure 5.**
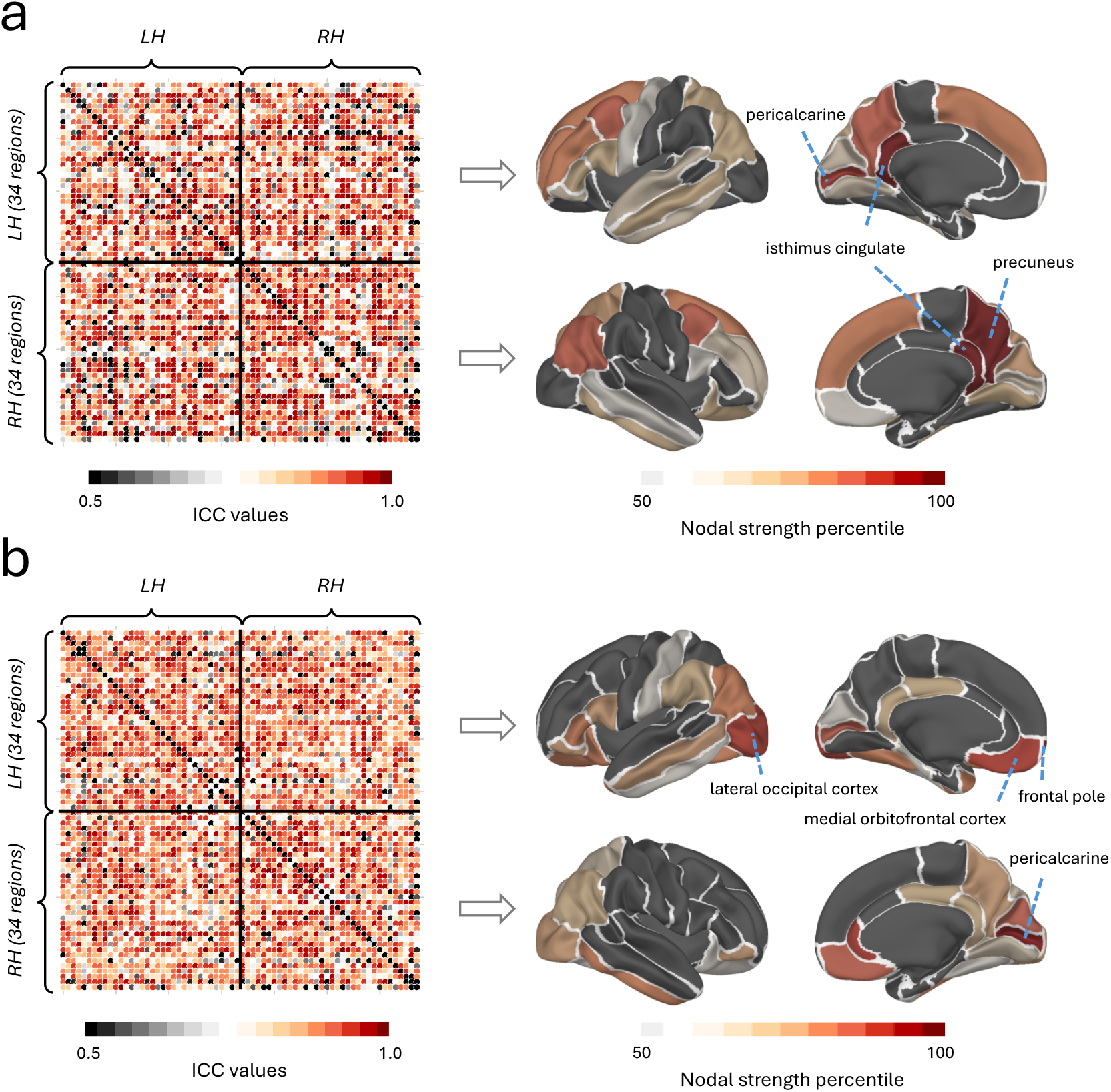
The spatial specificity of functional connectivity (FC) fingerprints in the tinnitus group was assessed using intra-class correlation (ICC), which quantified the fingerprint of each brain edge (connection) for the (**a**) alpha and (**b**) gamma frequency bands. Fingerprinting hubs were identified by calculating the nodal strength of the ICC matrix for each frequency band. Regions with nodal strength in the top 50th percentile were visualized on the cortical surface, highlighting the key hubs contributing to the altered connectivity patterns. In the alpha band (**a**), the highest nodal strength was observed in the left and right isthmus cingulate, precuneus (right hemisphere), and pericalcarine (left hemisphere). In the gamma band (**b**), the dominant hubs included the right pars orbitalis, right rostral middle frontal gyrus, and left superior frontal gyrus.

## 4. Discussion

In this study, we aimed to address several key questions:i) We evaluated the robustness of the graph learning method with respect to the number of samples, particularly investigating whether the brain functional connectome could be inferred reliably from short-duration recordings. ii) We analyzed group-level differences in functional connectivity (FC) between healthy individuals and those with tinnitus. iii) We explored whether individuals or groups could be accurately identified based solely on their unique patterns of brain activity. iv) Finally, we examined the spatial configuration of brain fingerprints within the tinnitus group to uncover distinct connectivity patterns.

To assess graph robustness, we examined whether consistent results emerged when using only a small fraction of time samples. This was achieved by comparing Portrait Divergence (PDiv) between these graphs and the one constructed from all data points. The PDiv comparison checks whether the global and local structures of the graphs are preserved, with smaller PDiv values indicating better preservation of these structures [48]. As PDiv is an Unknown Node-Correspondence (UNC) method and is a general and flexible metric capable of handling graphs with different numbers of nodes, a Known Node-Correspondence (KNC) method was also employed. In this method, the sum of the Euclidean distances between corresponding edges in the graphs and the optimal graph was computed. The analysis confirmed that the graphs remained stable regardless of the number of time samples used, across various parameter settings (Figure 1.b). The graph learning (GL) method was also compared with the PLI method. For any sample size, both methods produced approximately equal PDiv values. Notably, for smaller sample sizes, negative PDiv values indicated smaller graph distances in the GL method. Euclidean distance comparisons further demonstrated that the GL method consistently outperformed PLI, especially for smaller sample sizes (Figure 1.c, d). The stability of the GL method with few time samples makes it suitable for functional connectivity estimation in task-related data, where ERPs are divided into shorter epochs. In future studies, we aim to explore this application by averaging graphs across trials to represent overall connectivity. This may overcome limitations associated with traditional methods like PLI and Coherence, which rely on estimating cross-spectral density using Morlet waveforms and are sensitive to edge effects and time-frequency trade-offs in short signals. Similarly, the rapid convergence of the GL method highlights its potential for future use in fast real-time MEG applications, where conventional correlation- or spectral-based approaches often fail due to limited data; nonetheless, our finding that GL can produce stable and meaningful connectivity patterns from brief time segments serves as a proof-of-concept for its potential suitability in tracking temporal dynamics in near real-time—though demonstrating true real-time applicability still requires further validation and implementation within a dedicated real-time processing framework. This issue also arises in real-time applications of MEG signal processing: Traditional correlation- or spectral-based methods often fail with such limited data, but the rapid convergence of the GL method renders it ideal for fast real-time applications. However, while the GL approach captures relationships between brain regions in MEG recordings, it fails to account for how these connections evolve over time because the analysis treats each time sample as an independent observation. A promising future direction would be to integrate a graph spectral representation of the instantaneous MEG data, as proposed by Shuman et al. [49], on top of the learned networks, potentially in combination with techniques like the Fukunaga-Koontz transform [50], to jointly model spatial and temporal dynamics of brain connectivity. While such approaches offer a more dynamic analysis of how brain networks evolve over time, treating functional connectivity as static also provides advantages—simplifying the analysis, reducing computational complexity, and making the method more tractable for smaller datasets.

The whole brain connectivity analysis at the group level showed significant differences (p < 0.05) in theta, alpha, and gamma frequency band connectivity, but not in the delta and beta bands (see Supplementary Table 2). This highlights the importance of these frequency band connectivities in distinguishing control subjects from those with tinnitus, aligning with previous studies [9, 8]. Yet, unlike [9], resulting differences in the alpha networks in our study were both negative and positive: In memory-related networks (alpha connectivity between IPU, INS, PHC, and SFL, see Figure 2), as part of the Default Mode Network (DMN), connectivity in the tinnitus group become less pronounced whereas increased alpha connectivity can be observed in more auditory-related connections (alpha connectivity between LOF and TT, see Figure 2) pertaining to the large-scale Auditory Network (AUN) and potentially the Salience Network (SAN), given the involvement of the orbitofrontal cortex. Thus, decreased alpha connectivity in memory-related regions suggests DMN hypoactivity in tinnitus, aligning with findings of altered DMN functioning [51]. Conversely, increased alpha connectivity between LOF and TT indicates hyperactivation of the AUN and possibly the SAN, aligning with enhanced auditory cortex excitability in tinnitus [52]. Our findings of reduced theta band connectivity in tinnitus patients, specifically between the fusiform gyrus and banks of the superior temporal sulcus (right hemisphere), the cuneus and lateral occipital cortex (left hemisphere), and the paracentral lobule and precentral gyrus, suggest disruptions in large-scale brain networks linked to the Visual Network (VN; cuneus and lateral occipital cortex), Sensorimotor Network (SMN; paracentral lobule and precentral gyrus), and potentially to the AUN (fusiform gyrus and superior temporal sulcus). These findings partially corroborate Schlee et al.’s [53] observation of widespread abnormal functional connectivity in tinnitus, extending beyond auditory areas to include visual and sensorimotor regions. Results are also in line with Mohan et al.’s [54] significant theta differences in their phase-lagged coherence-based global graph connectivity strength analysis in source space. Methodological differences between our and the former analyses, namely the application of Partial Directed Coherence in the case of Schlee et al. [53] and the graph-theoretical analysis on lagged phase coherence-based connectivity matrices in Mohan et al. [54], are to be noted here. Yet, further adding to the sensitivity, specificity and validity of our method, results mostly overlap in presence of these differential methods and data sources.

The observed theta-band disruptions also resonate with findings from fMRI studies. Similar disruptions have been reported in fMRI studies, such as altered DMN and VN connectivity [55], though primarily in unilateral tinnitus (while our sample mostly consists of individuals with bilateral tinnitus, see Supplementary Table 1). This suggests that reduced theta connectivity in tinnitus reflects impaired integration across sensory and somatosensory networks. Finally, our findings of stronger gamma band connectivity in tinnitus patients between regions such as the inferior parietal cortex, precuneus, pars orbitalis, and superior temporal sulcus align with previous M/EEG studies. Increased gamma connectivity between frontal, parietal, and temporal regions has been reported, reflecting heightened attentional and sensory processing [53, 56]. Synthesizing our findings with previous studies, tinnitus is associated with increased gamma connectivity across auditory, attentional, and limbic networks, indicating maladaptive changes in sensory, attentional, and emotional integration. Taken together, our findings, combined with previous research, suggest that tinnitus involves complex alterations in functional connectivity across multiple frequency bands and large-scale brain networks, particularly affecting the integration of sensory, attentional, and emotional processing. Future studies should focus on characterizing the temporal dynamics of these connectivity patterns and their relationship to tinnitus severity, while exploring targeted neuromodulation techniques [57] to normalize altered connectivity patterns in specific networks and frequency bands.

Our study demonstrated that individuals can be accurately identified based solely on their brain activity patterns. Notably, individuals exhibited high consistency in their brain connectivity across test and retest sessions (*I*_*sel f*_), regardless of their clinical condition. Identification ranks for both groups (Fig. 3.b) indicated that graph learning (GL) enhances both global and local (edgewise) individual fingerprints of functional connectivity profiles. This improvement was data-driven and independent of group membership. Specifically, by “identification,” we refer to within-session identification, where two sample time frames from the same individual’s resting-state MEG are matched as belonging to the same subject, from a pool of samples across participants. This does not imply subject identification in an absolute or clinical sense, but rather demonstrates that the learned functional connectomes are stable and reproducible, and carry individual-specific signatures. It is worth noting that *I*_*di f f*_, which captures the difference between intra- and inter-subject similarity, is computed across the entire dataset. While the inter-subject component can in principle be influenced by group composition, our study used a fixed cohort throughout. Thus, the relative differences in *I*_*di f f*_ across individuals remain meaningful within this stable context. Another potential concern is that splitting the resting-state recording into two halves for the same individual (test and retest) might introduce a risk of circularity as they might share similar noise covariance structures and potentially inflating the results. To address this, we compared GL ranks with those from a correlation-based method (PLI). The GL method outperformed PLI, achieving more accurate individual identification (Fig. 3.b). This also suggests that the identifiability achieved by GL is not solely driven by shared noise but rather reflects meaningful and stable subject-specific connectivity features. Although specific connections distinguishing healthy individuals from those with tinnitus were identified, substantial variability across subjects highlights the need for personalized analyses in future studies. Spectral clustering on the similarity matrix from the identifiability matrix (see Section 3 for details) visualized with 2D MDS, revealed significant variability in subjects’ functional connectivities (FCs). Notably, individuals appearing isolated in the MDS representation were typically those with high *I*_*di f f*_ values (Fig. 4.a), which may artificially inflate the silhouette score—an index of clustering quality—particularly when partitioning the data into two clusters. Consequently, the three-cluster solution likely reflects a combination of true underlying substructure and methodological effects driven by inter-individual differences in identifiability. While the silhouette scores for both the two- and three-cluster solutions were relatively low (0.21 and 0.202, respectively), this is expected in data with high individual variability and weakly separated clusters. Our goal was not to claim strong categorical separability, but rather to explore and illustrate patterns of variability in FC profiles. Notably, individuals with high *I*_*di f f*_ values tend to appear as outliers in the similarity space, which contributes to lower cohesion and separation as captured by the silhouette metric. These patterns, even if subtle from a clustering validation standpoint, provide valuable insight into the heterogeneity of FCs among individuals. Furthermore, these findings underscore the importance of focusing on individual variability rather than solely on group differences when studying functional connectivity alterations in tinnitus. We further demonstrated that the GL method effectively captures the underlying functional connectome, differentiating individuals while maintaining group coherence. In contrast, PLI scattered individuals without forming clear group structures (Fig. 4.b), highlighting the superior performance of the GL method in preserving individual and group-level distinctions. While the GL method allows for FC estimation from relatively small sample sizes by leveraging signal smoothness priors, it is important to acknowledge that this flexibility does not eliminate the fundamental need for sufficient statistical power. In particular, efforts to characterize inter-individual variability—a central goal of personalized neuroscience—require a sample large enough to robustly estimate population-level variability and distinguish it from measurement noise or artifacts. The high dimensionality of FC matrices, coupled with potential variability in signal-to-noise ratio across individuals, compounds the difficulty of reliable estimation in small cohorts. In our current sample, we are cautious in interpreting fine-grained individual differences, and we view our findings as preliminary evidence motivating larger-scale follow-up studies. Another important consideration is the need for methods that better capture individual-level deviations, beyond traditional case-control comparisons. Approaches like normative modeling [58] have emerged as powerful tools to address this challenge by establishing a model of typical brain function (e.g., in healthy controls) and quantifying how individual subjects deviate from this norm. These models can identify atypical brain patterns without relying solely on group membership, enabling a more nuanced understanding of disorder-related changes and potentially revealing meaningful subgroups or phenotypes within heterogeneous populations. While our current study was not designed to implement normative modeling, future work could incorporate such methods to more explicitly characterize individual variability and relate deviations from normative patterns to clinical outcomes.

Examining the topological distribution of connections underlying this unique pattern, we observed a spatial reconfiguration of regions with the highest idiosyncratic fingerprint within tinnitus group (Fig. 5.a, b left panels). These regions may, in fact, represent areas least affected by tinnitus, as they remain distinct across individuals rather than converging toward a common pattern. To identify regions impacted by tinnitus, we compared their spatial specificity to that of controls (Supplementary Figure S1). By examining regions with heightened spatial node strength in the tinnitus group, we identified key areas that play a more significant role in distinguishing individuals with tinnitus. In the alpha frequency band, these include the left and right isthmus of the (posterior) cingulate gyrus, while in the gamma band, prominent regions include the left frontal pole, the right pericalcarine cortex, and the left medial orbitofrontal cortex. In these cortical regions, connectivity profiles exhibit subject-specific idiosyncrasies that extend beyond the variability seen in controls. Conversely, regions with greater spatial node strength in controls appear to have diminished discriminative power for tinnitus, exhibiting more homogeneous connectivity patterns in affected individuals. These regions include the left and right superior frontal cortex, right temporal pole, and right rostral anterior cingulate cortex in the alpha band, along with the right pars orbitalis, right rostral middle frontal cortex, and left superior frontal cortex in the gamma band. This homogenization may reflect tinnitus’ impact on large-scale brain network organization.

The lack of open-access MEG datasets for tinnitus populations poses a significant challenge to independent validation and replication of our findings. Unfortunately, we could not get hold of another tinnitus MEG data set at this point. Furthermore, given the small sample size in our study, it was not feasible to divide the data into separate training and validation cohorts. However, we recognize the importance of validating our results in an independent cohort, and this will be an important direction for future research. Future work could also benefit from methodological extensions—for example, incorporating priors from structural connectivity, developing hybrid supervised–unsupervised graph estimation methods, or optimizing graph learning approaches for clinical outcome prediction. However, the key contribution of this work lies in the novel approach to extracting individual features within each specific diagnostic group. While traditional machine learning methods focus on extracting features to allocate new patients to specific diagnostic categories, our approach emphasizes identifying sources of individual variability unique to each group. This can be instrumental in predicting clinical features at an individual level within the diagnostic group. In this context, our fingerprinting approach, which enables the extraction of individualized functional connectivity features, holds potential for use in machine learning-based predictions. Importantly, diagnostic classification and individualized feature extraction—are complementary, addressing distinct clinical and research questions. Together, they offer a more comprehensive framework for understanding and leveraging brain connectivity in clinical settings.

## Conclusion

In this study, we present a graph learning approach to extract frequency-specific functional networks from MEG data. Our results demonstrate that the obtained networks outperform conventional correlation-based ones in terms of fingerprinting accuracy. At the group level, our study reveals previously unreported patterns of altered connectivity in tinnitus patients across multiple frequency bands, involving canonical functional networks DMN, AUN, VN, and SAN. Globally, we observed reduced theta-band connectivity in regions associated with the VN and SMN, decreased alpha connectivity in memory-related DMN regions coupled with increased alpha connectivity in auditory-related areas, and increased gamma-band connectivity between auditory, attentional, and limbic networks, suggesting a complex reorganization of large-scale brain networks in tinnitus that extends beyond auditory areas. In more detail, these specific alterations across the theta, alpha, and gamma bands constitute key novel findings stemming from our pioneering application of GL to MEG data to investigate tinnitus-related functional connectivity. Our GL-MEG approach particularly identified previously uncharacterized details such as: significantly reduced theta-band connectivity—specifically between the fusiform gyrus and banks of the superior temporal sulcus (right hemisphere), which may reflect impaired integration of auditory information and reduced long-distance communication within the AUN and with broader brain networks, as well as between the cuneus and the lateral occipital cortex (left hemisphere), and between the paracentral lobule and the precentral gyrus—indicative of disruptions across VN and SMN in addition to these AUN alterations; a complex pattern of alpha-band alterations, featuring decreased connectivity in memory-related DMN regions (e.g., between IPU, INS, PHC, and SFL) alongside increased connectivity in auditory-related pathways (e.g., between LOF and TT) implicating the AUN and SAN, where such increased alpha connectivity within auditory cortices could signify a more rigidly coupled, internally focused processing state or an altered inhibitory gating mechanism related to the tinnitus percept; and stronger gamma-band connectivity between regions such as the inferior parietal cortex, precuneus, pars orbitalis, and superior temporal sulcus, suggesting increased integration across AUN, attentional, and limbic networks. In addition, we investigated the spatial specificity of functional network fingerprints by analyzing the distinctiveness of each connection at the individual level. This analysis uncovered key brain regions associated with tinnitus, most importantly key DMN nodes in prefrontal and parietal/occipital regions. Our study provides novel insights into the neural mechanisms underlying the condition. Finally, the proposed approach offers rapid and reliable implementation, paving the way for objective diagnostics and personalized treatment strategies, including real-time interventions.

## Data availability

Large raw data files, including resting state MEG recordings and MR images are available upon request. As data is not deidentified, data access will require a data processing agreement and ethics approval.

## Code availability

The codes used for analyzing MEG and MRI data, graph learning and generating figures are available in the following repository: https://github.com/payamsash/paypatmeg.

## Conflict of Interest Disclosures

C.R.C. reported being a member of the British Tinnitus Association’s Professional Advisers’ Committee and the American Tinnitus Association’s Scientific Advisory Board.

## Acknowledgments

P.S.S. and P.N. have been funded by SNF (Project Grant number 325130 208164) for this work. C.R.C. received funding from the Rainwater Charitable Foundation, Tysta Skolan, the Swedish Research Council (VR 2023-02326), the GENDER-Net Co-Plus Fund (GNP-182), the European Union’s Horizon 2020 Research and Innovation Programme, grant agreement no. 848261, and the European Union’s Horizon 2020 research and innovation program under Marie Skłodowska-Curie grant agreement no. 722046, and a Karolinska Institutet Doctoral (KID) grant. N.K.E. received funds from Hörselforskningsfonden and Tysta Skolan.

## Author contributions

P.S.S. analyzed the data, generated figures and wrote the first draft of the paper and all authors contributed to and approved the final manuscript. H.H.B. supervised, reviewed and edited the graph analysis sections. D.V.D.V. and T.K reviewed and edited the manuscript. N.K.E. and C.R.C. collected and shared raw MEG + MRI data and P.N. supervised, reviewed and edited the neurophysiological relevance of the results.

## Notes

### Competing Interest Statement

Christopher R. Cederroth reported being a member of the British Tinnitus Associations Professional Advisers Committee and the American Tinnitus Associations Scientific Advisory Board.

### Summary of Updates

Some updates on Figures and extra explanation on the text.

