## Supplementary information for "Frequency-Specific Resting-State MEG Network Characteristics of Tinnitus Patients Revealed by Graph Learning"

*Comparison of head movement between the two groups*

We computed framewise displacement (FD) for each subject using the continuous head position indicator (cHPI) traces recorded during MEG acquisition. Specifically, FD at time t was calculated as:

$$\mathrm{FD}_t = |\Delta x_t| + |\Delta y_t| + |\Delta z_t| + r \cdot |\theta_t|$$

which represents the frame-to-frame displacement along the 3 axes. $r$ is radius of the head sphere (commonly assumed to be 50 mm) and $\theta_t$ is the rotation angle between consecutive head position matrices, computed as:

$$\theta_t = \cos^{-1}\left(\frac{\mathrm{Tr}(R_{t-1}^{\top}R_t) - 1}{2}\right)$$

A two-sample t-test comparing mean FD between tinnitus patients and control subjects yielded: $t(39) = -0.707, p = 0.483$, indicating no significant difference in head motion between the groups. Furthermore, we would like to emphasize that the MEGIN MaxFilter algorithm already incorporates movement compensation using the cHPI data by realigning the signals to a common head position and removing external interference through Signal Space Separation (SSS). These steps help mitigate potential motion-related confounds in downstream connectivity analysis. Together, these analyses suggest that residual head motion is unlikely to explain the observed group differences in functional connectivity.

Table 1: Descriptive characteristics of the participants. Between parenthesis are either standard deviation (SD) or percentage (%). Abbreviations: Perceived Stress Questionnaire (PSQ), Hyperacusis Questionnaire (HQ), Hospital Anxiety and Depression Score (HADS), Pure Tone Audiometry (PTA), High Frequency (HF), decibel hearing levels (dB HL)

| | **Control group (n=23)** | **Tinnitus group (n=18)** |
|---|---|---|
| **Age** | | |
| Mean (SD) | 29.17 (5.69) | 36.83 (7.66) |
| **Sex** | | |
| Male | 11 (47.8%) | 11 (61.1%) |
| Female | 11 (47.8%) | 7 (38.9%) |
| Other | 1 (4.3%) | 0 (0.0%) |
| **Handedness** | | |
| Right | 21 (91.3%) | 17 (94.4%) |
| Left | 2 (8.7%) | 1 (5.6%) |
| **Tinnitus** | | |
| Yes, always | 0 (0.0%) | 18 (100.0%) |
| Yes, often | 0 (0.0%) | 0 (0.0%) |
| Yes, sometimes | 0 (0.0%) | 0 (0.0%) |
| Not last year | 6 (26.1%) | 0 (0.0%) |
| No, never | 16 (69.6%) | 0 (0.0%) |
| Don't know | 1 (4.3%) | 0 (0.0%) |
| **Tinnitus lateralization** | | |
| Right | - | 1 (5.6%) |
| Left | - | 5 (27.8%) |
| Both | - | 12 (66.7%) |
| **PSQ score** | | |
| Mean (SD) | 0.23 (0.14) | 0.41 (0.15) |
| **HQ score** | | |
| Mean (SD) | 9.70 (6.38) | 22.61 (8.23) |
| **HADS Anxiety score** | | |
| Mean (SD) | 4.09 (3.12) | 8.61 (3.71) |
| **HADS Depression score** | | |
| Mean (SD) | 1.91 (2.63) | 4.78 (4.11) |
| **THI score** | | |
| Mean (SD) | - | 43.78 (27.56) |
| **PTA Left (dBHL)** | | |
| Mean (SD) | 3.00 (2.91) | 6.85 (6.12) |
| **HF PTA Left (dBHL)** | | |
| Mean (SD) | 8.65 (12.51) | 25.59 (22.72) |
| **PTA Right (dBHL)** | | |
| Mean (SD) | 3.87 (3.09) | 6.93 (6.44) |
| **HF PTA Right (dBHL)** | | |
| Mean (SD) | 8.54 (9.09) | 22.68 (20.80) |
| **Tin pitch (kHz)** | | |
| Mean (SD) | - | 9.94 (7.44) |
| **Tin loudness (dBHL)** | | |
| Mean (SD) | - | 24.35 (21.73) |
| **Tin loudness (dBSL)** | | |
| Mean (SD) | - | 3.38 (8.82) |

Table 2: The effect sizes (Hedges' g) for all connections that showed significant differences between the tinnitus and control groups within specific frequency bands are presented below.

| Region 1 | Region 2 | Frequency band | Effect size |
|---|---|---|---|
| bankssts-rh | fusiform-rh | theta | 1.4365 |
| cuneus-lh | lateraloccipital-lh | theta | 1.4071 |
| paracentral-rh | precentral-lh | theta | 1.3165 |
| inferiorparietal-lh | precuneus-lh | alpha | 1.2714 |
| insula-lh | parahippocampal-lh | alpha | 1.4618 |
| lateralorbitofrontal-lh | transversetemporal-lh | alpha | -1.1363 |
| parahippocampal-lh | superiorfrontal-lh | alpha | -1.6720 |
| parahippocampal-lh | superiorfrontal-rh | alpha | -1.6627 |
| bankssts-lh | parsorbitalis-lh | gamma | -1.8622 |
| frontalpole-lh | superiorfrontal-lh | gamma | 1.4645 |
| frontalpole-rh | superiorfrontal-lh | gamma | 1.5075 |
| frontalpole-rh | superiorfrontal-rh | gamma | 1.7006 |
| inferiorparietal-lh | parsorbitalis-lh | gamma | -1.4549 |
| inferiorparietal-lh | precuneus-lh | gamma | 1.2350 |
| paracentral-lh | superiorparietal-lh | gamma | 1.0708 |

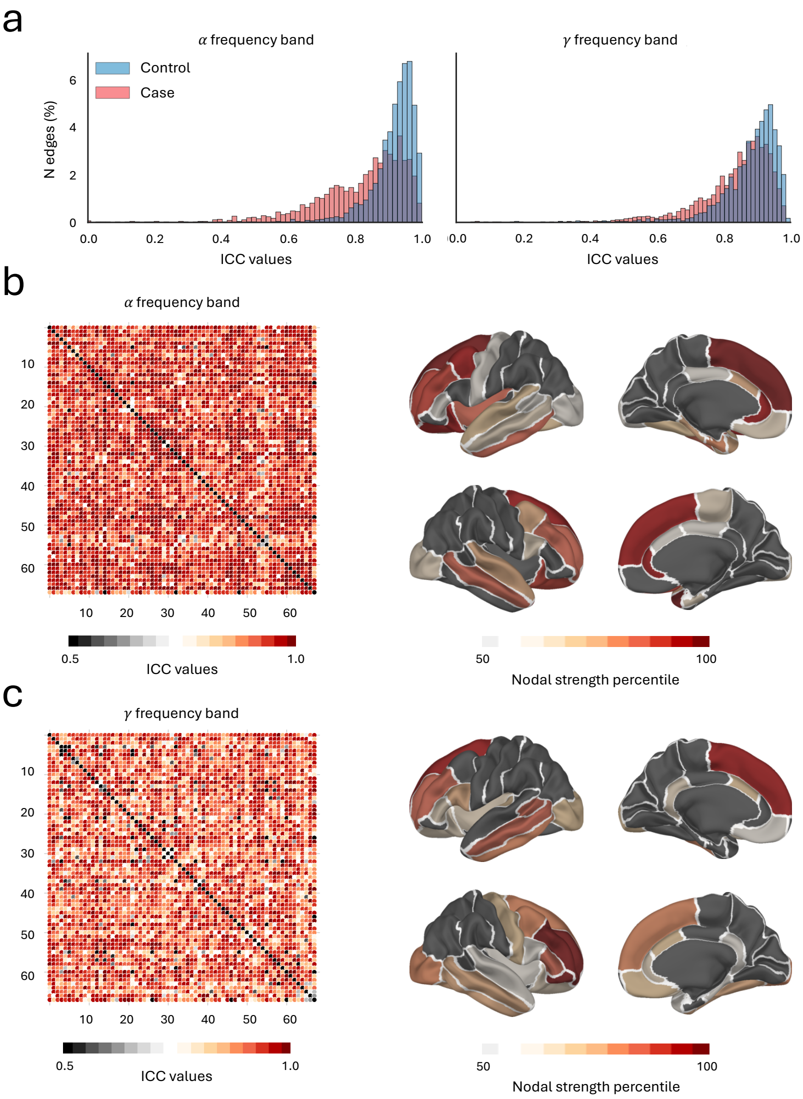


Figure S1: The spatial specificity of functional connectivity (FC) fingerprints within the control group was evaluated using intra-class correlation (ICC), which measured the importance of each connection in the alpha and gamma frequency bands in discriminating individuals. (**a**) The ICC value distributions for both groups, shown for the alpha band (left panel) and gamma band (right panel), reveal predominantly lower ICC values in the tinnitus group. This pattern suggests greater similarity between functional connectivity within the tinnitus group. (**b**) The nodal strength of the ICC matrix was calculated for each frequency band. Brain regions with nodal strength in the top 25th percentile were mapped onto the cortical surface, revealing the key hubs that contribute to the altered connectivity patterns observed in the control group.

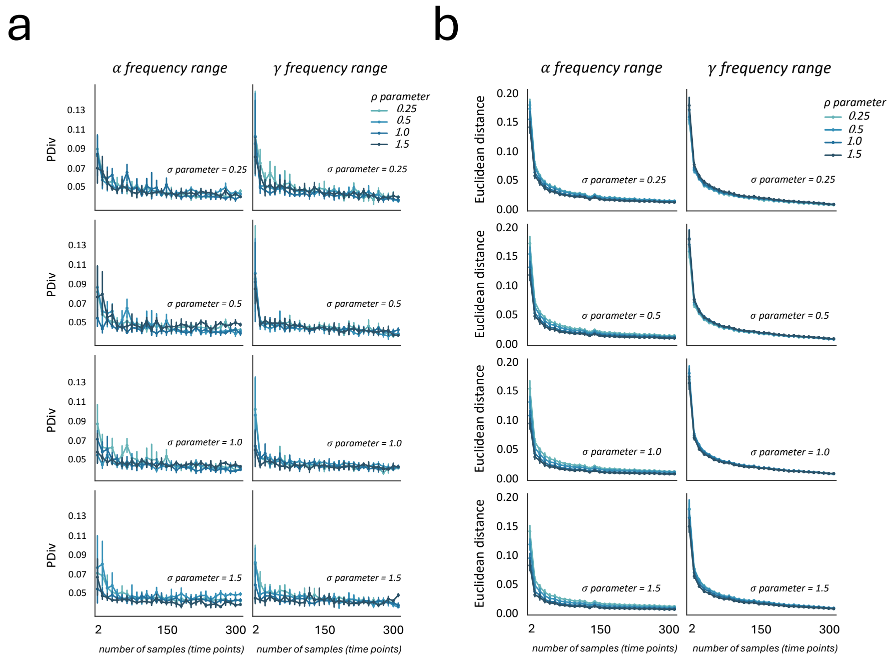


Figure S2: Graph stability across various regularization parameter configurations and different numbers of time points used for graph learning; PDiv (**a**) and Euclidean distances (**b**) were calculated between each graph and the ideal graph at alpha and gamma frequency ranges.

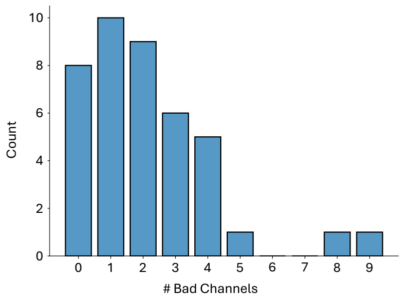


Figure S3: Histogram showing the number of MEG channels identified as bad (noisy or flat) per recording. Bad channels were automatically annotated and excluded from further analysis. The MEG system includes 102 magnetometers and 204 gradiometers in total.

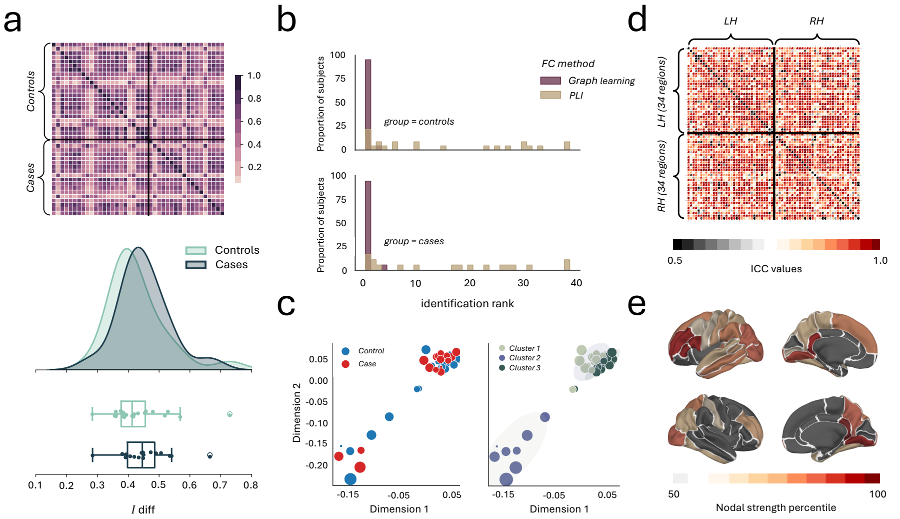


Figure S4: (**a**) The identifiability matrix shows within-subjects similarity (Iself, diagonal elements) and between-subjects similarity (off-diagonal elements) across all subjects (tinnitus + control) for theta frequency band. The density and box plots of Idiff values are shown below. The Idiff value for each subject is calculated as the difference between their Iself value and the average of their off-diagonal values, highlighting the subject's distinctiveness in functional connectivity. (**b**) Bar plots showing the identification ranks in two groups (tinnitus and control) within theta frequency band derived via the GL and PLI methods, comparing the test and retest FCs among subjects. (**c**) The 2D MDS representation of the similarity matrix, derived from the identifiability matrix for the theta frequency band, shows a clear grouping of controls and tinnitus patients (left panel) together with the clustering results (right panel) (**d**) The spatial specificity of functional connectivity (FC) fingerprints in the tinnitus group was assessed using intra-class correlation (ICC), which quantified the fingerprint of each brain edge (connection) for the theta frequency band. (**e**) Fingerprinting hubs were identified by calculating the nodal strength of the ICC matrix for each frequency band. Regions with nodal strength in the top 50th percentile were visualized on the cortical surface, highlighting the key hubs contributing to the altered connectivity patterns.

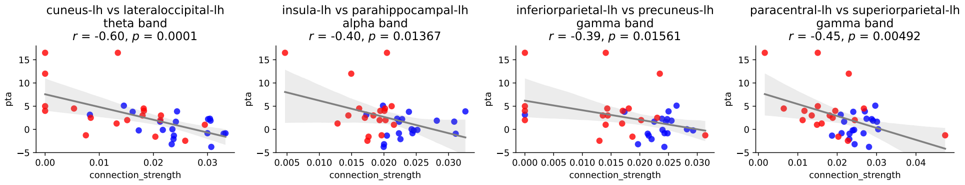


Figure S5: Correlations between Pure Tone Average (PTA) and the connections that showed statistically significant differences between the two groups are presented. Only connections with significant correlations are shown. Blue dots represent control subjects, and red dots represent individuals with tinnitus.

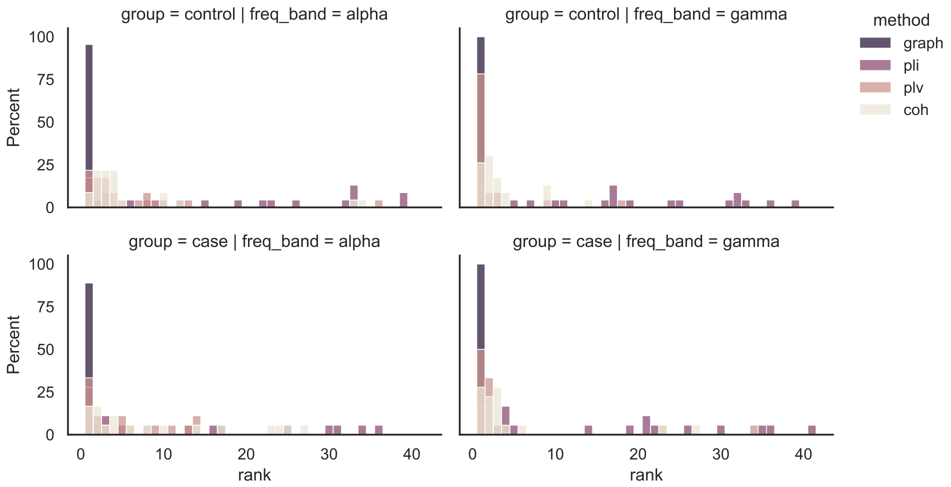


Figure S6: Bar plots show the identification ranks for the tinnitus and control groups using four different methods: Graph Learning (GL), Phase Lag Index (PLI), Phase Locking Value (PLV), and Coherence (Coh). The comparison is based on test–retest functional connectivity across subjects. It is evident that the GL method outperforms the other three approaches.
